# Acarbose suppresses symptoms of mitochondrial disease in a mouse model of Leigh Syndrome

**DOI:** 10.1101/2022.01.31.478591

**Authors:** Alessandro Bitto, Anthony S. Grillo, Ian B. Stanaway, Bao M. G. Nguyen, Kejun Ying, Herman Tung, Kaleb Smith, Ngoc Tran, Gunnar Velikanje, Silvan R. Urfer, Jessica M. Snyder, Ernst-Bernhard Kayser, Lu Wang, Daniel L. Smith, J. Will Thompson, Laura DuBois, William DePaolo, Matt Kaeberlein

## Abstract

Mitochondrial diseases represent a spectrum of disorders caused by impaired mitochondrial function ranging in severity from mortality during infancy to progressive adult-onset disease. Mitochondrial dysfunction is also recognized as a molecular hallmark of the biological aging process. Rapamycin, a drug that increases lifespan and health during normative aging also increases survival and reduces neurological symptoms in a mouse model of the severe mitochondrial disease Leigh Syndrome. The Ndufs4 knockout (*Ndufs4^-/-^*) mouse lacks the complex I subunit NDUFS4 and shows rapid onset and progression of neurodegeneration mimicking patients with Leigh Syndrome. Here we show that another drug that extends lifespan and delays normative aging in mice, acarbose, also suppresses symptoms of disease and improves survival of *Ndufs4^-/-^* mice. Unlike rapamycin, acarbose rescues disease phenotypes independently of mTOR inhibition. Furthermore, rapamycin and acarbose have additive effects in delaying neurological symptoms and increasing maximum lifespan in *Ndufs4^-/-^* mice. We find that acarbose remodels the intestinal microbiome and alters the production of short chain fatty acids. Supplementation with tributyrin, a source of butyric acid, recapitulates some effects of acarbose on lifespan and disease progression. This study provides the first evidence that alteration of the gut microbiome may impact severe mitochondrial disease and provides further support for the model that biological aging and severe mitochondrial disorders share underlying common mechanisms.

## Introduction

Mitochondrial disease is a term encompassing numerous disorders characterized by loss of function in one or more mitochondrial processes ^1,2^. These mitochondrial dysfunctions affect primarily energy-demanding tissues, such as skeletal muscle, heart, and brain, and manifest with different degrees of severity as seizures, myopathies, cardiomyopathies, encephalopathies, and cerebellar ataxias. Currently, no known cure or treatment is available for any of these pathologies, and it is estimated that at least 1 in 5000 live births suffer from severe forms of mitochondrial disease, with crippling consequences for quality of life and extremely reduced life expectancy ^3^.

Mutations in Complex I (NADH:Ubiquinone Oxidoreductase) of the mitochondrial electron transport chain have been reported in up to 30% of pediatric mitochondrial diseases, including severe pathologies such as Mitochondrial Encephalomyopathy with Lactic Acidosis and Stroke-Like episodes (MELAS), Fatal Infantile Lactic Acidosis, Leukoencephalopathy, Neonatal Cardiomyopathy, and Leigh Syndrome, a neurometabolic, degenerative disorder which is often fatal in the first three years of life ^1,4^. Mutations in the structural subunit of Complex I NADH-Dehydrogenase Ubiquinone Fe-S Protein 4 (NDUFS4) often cause loss of function of the whole complex and have been associated with severe forms of Leigh Syndrome ^5^. Knock out of exon 2 of the *Ndufs4* gene in mouse (*Ndufs4^-/-^*) recapitulates several aspects of Complex I deficiencies, including encephalopathy, retarded growth rate, lethargy, loss of motor skills, lactic acidosis, and extremely reduced lifespan ^6^.

Daily administration of rapamycin, an inhibitor of the nutrient sensing mechanistic target of rapamycin complex I (mTORC1), rescues multiple manifestations of mitochondrial disease in *Ndufs4^-/-^* mice, including neurological symptoms, accumulation of glycolytic intermediates and lactate, loss of body fat, aberrant nutrient and growth factor signaling, and survival ^7,8^. Hyperactivation of mTORC1 has been observed in brain of *Ndufs4^-/-^* mice as well as other models of mitochondrial disease ^7,9,10^; however, the precise mechanisms by which rapamycin attenuates mitochondrial dysfunction remains unknown. A metabolic shift away from glycolysis may partially contribute to disease rescue; prior evidence suggests this may be mediated in part through mTORC1 signaling in the liver may contribute ^11^ to disease rescue, along with inhibition of protein kinase C (PKC) signaling in the brain ^12^. Importantly, there is initial data indicating that rapamycin therapy may improve health in patients suffering from adult onset MELAS syndrome ^13^ as well as pediatric Leigh Syndrome ^14^.

Inhibition of mTORC1 with rapamycin is also a robust intervention for slowing or reversing biological aging. Rapamycin treatment improves health and increases lifespan during normative aging across multiple organisms including yeast ^15^, worms ^16^, fruit flies ^17^, and mice ^18^. In mice, lifespan extension has been reported in multiple strain backgrounds, across a broad dose range involving both dietary feeding and intraperitoneal injection ^18–28^. Age-related phenotypes where rapamycin has been reported to have positive effects include lower incidence of age-related cancers ^19,29^, protection against age-related cognitive dysfunction ^30,31^, preservation of tendon ^32^, improved heart function ^33–35^, restoration of immune function ^36^, improved kidney function ^37^, rejuvenation of oral health ^38,39^, improved intestinal function and reduced gut dysbiosis ^27,40^, and preservation of ovarian function ^41^. Although early, there is initial evidence suggesting similar effects on aging heart in dogs ^42^ and in immune function in humans ^43,44^.

Mitochondrial dysfunction is one of the primary hallmarks of aging ^45^ which has been implicated in numerous age-related disorders and functional declines ^46–48^. While it is clear that severe mitochondrial disease does clinically resemble accelerated aging, this connection has led us to hypothesize that aging is, to some extent, an acquired mitochondrial disease ^49^. The observation that mTOR dysregulation is shared between mitochondrial disease and normative aging can be viewed as support for this hypothesis, as can the fact that rapamycin is among the most effective known interventions at alleviating aspects of both conditions.

In order to further test the relationship between mitochondrial disease and normative aging, we set out to test whether the drug acarbose could impact disease progression and survival in *Ndufs4^-/-^* mice. Acarbose is an inhibitor of intestinal alpha-glucosidases and pancreatic alpha-amylases approved for the management of non-insulin dependent diabetes ^50^. Similar to rapamycin, acarbose has also been found to improve health and lifespan in mice ^51,52^. Here we report the discovery that acarbose has comparable efficacy to rapamycin at delaying disease progression and increasing survival in *Ndufs4^-/-^* mice. Unlike rapamycin, acarbose treatment does not inhibit mTOR signaling but instead remodels the intestinal microbiome to produce higher levels of the short-chain fatty acid butyrate. Butyrate supplementation is sufficient to partially rescue disease phenotypes, suggesting that acarbose may ameliorate the progression of mitochondrial disease by altering the composition of the intestinal microbiome. Altogether, these results provide further support for the hypothesis that aging and mitochondrial disorders share common underlying mechanisms and introduce the microbiome as a potential area of intervention to treat severe mitochondrial disease.

## Results

### Acarbose rescues disease phenotypes and improves survival in *Ndufs4^-/-^*mice

*Ndufs4^-/-^* mice older than post-natal day (p.n.) 38 present characteristic brain lesions, including vacuolation in the vestibular nuclei, cerebellar vermis, and olfactory bulb; increased vascularity of the brainstem and posterior cerebellum; and widespread astroglial and microglial reactivity ^6,53,54^. In order to determine whether acarbose can mitigate the appearance of these lesions, we fed *Ndufs4^-/-^* mice with either regular chow or chow containing 1000 ppm acarbose from weaning (around post-natal day 21) to 50 days of age, when mice were sacrificed. Mice were anesthetized with a ketamine/xylazine mix and perfused intracardially with phosphate buffered saline (PBS) and 10% neutral buffered formalin. Deterioration of mitochondrial function is more prominent in the olfactory bulb, brainstem, and cerebellum of *Ndufs4^-/-^* brains ^54^. To better determine whether acarbose rescues signs of pathology in these regions, we sectioned 10% neutral-buffered formalin perfused and fixed brains. We examined the levels of brain astrogliosis and microgliosis in the brains of *Ndufs4^-/-^* mice and in mice treated with acarbose by performing immunohistochemistry against Glial Fibrillary Acidic Protein (GFAP), a marker of astrocyte activation, and Ionized calcium-binding adapter molecule 1 (Iba1), a marker of microglia activation. As expected, *Ndufs4^-/-^* mice displayed abundant GFAP and Iba1 staining across the brain, with particularly high scores in the most affected regions (olfactory bulb, brainstem/cerebellum.) Acarbose significantly reduced GFAP staining in the olfactory bulb (section 1,) medulla oblongata, and cerebellum (section 5) (Figure 1B, and C,) and showed a trend towards reduced Iba1 staining in the same sections of *Ndufs4^-/-^* mice (Figure S1A and B). Acarbose treatment was also associated with a trend toward reduced levels of vacuolation in the vestibular nuclei that did not reach statistical significance (Figure S1C).

**Figure 1:**
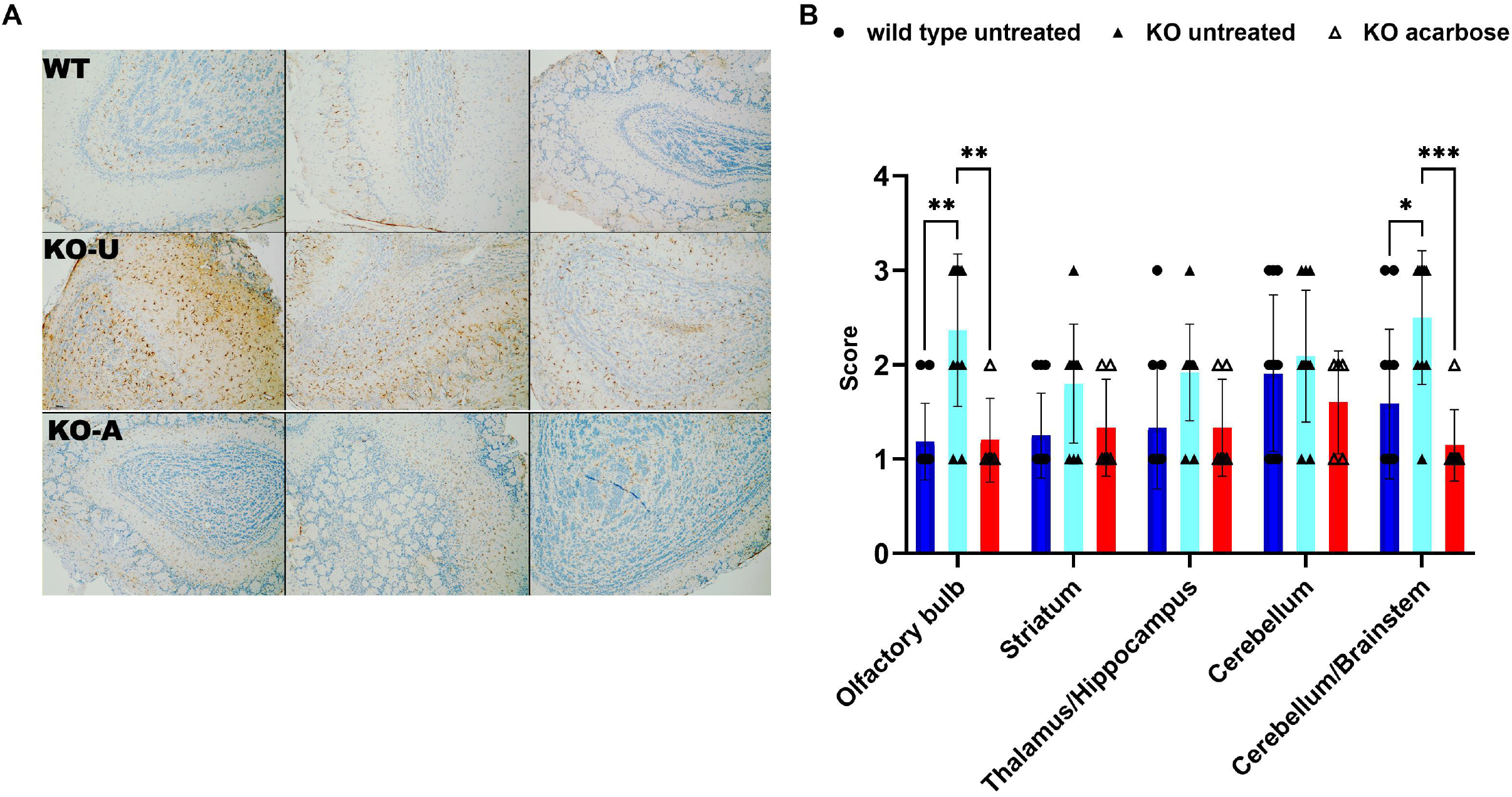
Acarbose reduces brain inflammation across the brain in *Ndufs4^-/-^* mice. **A.** Representative images of GFAP immunostaining in the olfactory bulb of wild type untreated, *Ndufs4^-/-^* untreated, and *Ndufs4^-/-^* mice treated with acarbose at post-natal day 50. **B.** Semi-quantitative scores of intensity of GFAP immunostaining in the sections described in panel A. **: p<0.01, *: p<0.05 vs untreated *Ndufs4^-/-^* mixed effects analysis, N=4-5/group.

Next, we sought to determine whether acarbose could suppress neurological phenotypes displayed by *Ndufs4^-/-^* mice. We observed mice treated with acarbose daily from post-natal day 21 until death or euthanasia and recorded the initial onset of hind-limb clasping, a characteristic neurological symptom of *Ndufs4^-/-^* mice ^6^. Consistent with the reduction in brain pathology (Figure 1) acarbose significantly delayed the onset of clasping by approximately 10 days or 23% (Figure 2A). Untreated *Ndufs4^-/-^* mice are growth-impaired and weigh consistently less than wild type or heterozygous littermates throughout their lifespan (Figure 2B and C). Additionally, they begin losing weight around 35 days post-natal as neurological symptoms begin to appear (Figure 2C). Both wild type and *Ndufs4^-/-^* mice treated with acarbose showed delayed growth compared to their isogenic controls (Figures 2B and C). *Ndufs4^-/-^* mice treated with acarbose reached their peak weight around 10 days later than untreated mice (p.n. 47 vs p.n. 37, Figure 2C). Consistent with the delay in the onset of clasping (Figure 2A,) acarbose-treated animals did not show a similar sharp decline in weight after onset of the disease (Figure 2C.) Lastly, we measured the effect of acarbose on survival and observed a 65% increase in median and maximum survival (Figure 2D, pooled results, Figure S2A, B for individual cohorts). In HET3 mice, acarbose extends lifespan during normative aging in males to a greater extent than female animals ^52,55^. However, we found no appreciable sex-specific difference in lifespan for acarbose-fed *Ndufs4^-/-^* mice (Figure S2C). Altogether, these results indicate that acarbose successfully treats symptoms of mitochondrial disease in *Ndufs4^-/-^* mice.

**Figure 2:**
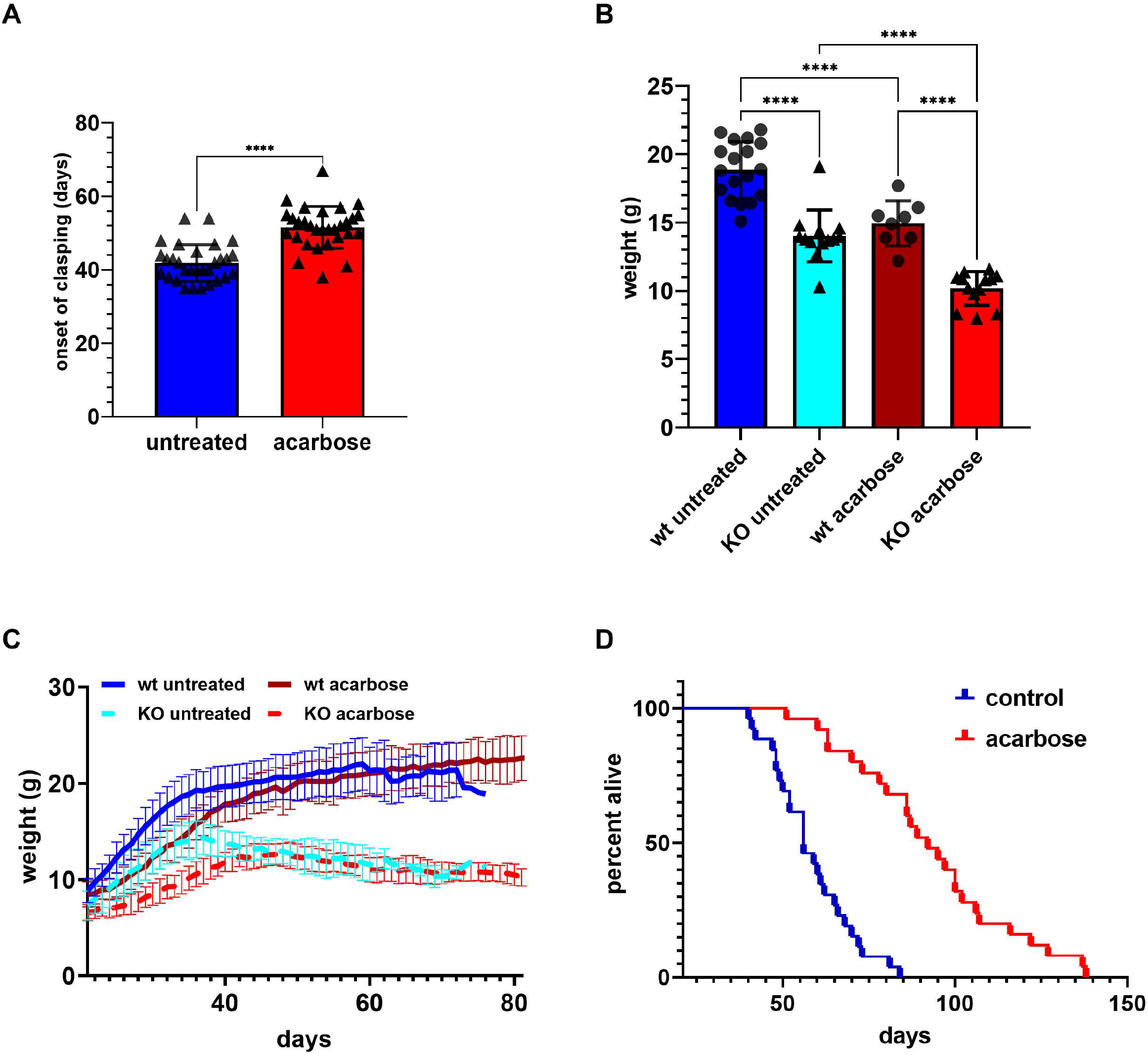
Acarbose increases survival and delays onset of disease in Ndufs4-/- mice. Wild type and *Ndufs4^-/-^* mice were fed either control chow or chow supplemented with 0.1% (1000 ppm) acarbose from weaning (post-natal day 21.) **A.** Onset of neurological symptoms (clasping) measured in days after birth in control-chow (blue) and acarbose-chow (red) fed *Ndufs4^-/-^* mice. **** p <0.0001 Student’s t test. N=30 **B.** Comparison of weights at 35 days post-natal for control-chow fed wild type, *Ndufs4^-/-^* mice, acarbose-chow fed wild type, and *Ndufs4^-/-^* mice. **** p<0.0001 one way ANOVA. N=18 wild type untreated, N=13 *Ndufs4^-/-^* untreated, N=8 wild type acarbose, N=15 *Ndufs4^-/-^* acarbose **C.** Weight progression from weaning until post-natal day 81. Solid dark blue: wild type mice fed control chow. Solid dark red: wild type mice fed acarbose chow. Dotted light blue: *Ndufs4^-/-^* mice fed control chow. Dotted light red: *Ndufs4^-/-^* mice fed acarbose chow. N=18 wild type untreated, N=13 *Ndufs4^-/-^* untreated, N=8 wild type acarbose, N=15 *Ndufs4^-/-^* acarbose **D.** Survival curves of *Ndufs4^-/-^* mice fed either control (blue) or 0.1% acarbose diet (red.) Median lifespan was 56 days for control-chow fed, and 92 days for acarbose-chow fed mice, log-rank p <0.0001. Data are pooled from two independent experiments (see figure S2.) N=26 untreated, N=25 acarbose-treated.

### Acarbose and rapamycin rescue mitochondrial disease by independent mechanisms

We investigated the putative mechanisms by which acarbose rescues disease in *Ndufs4^-/-^* mice. Acarbose is an inhibitor of alpha-glucosidases and alpha-amylases that effectively reduces absorption of carbohydrates through the intestinal mucosa ^50,56,57^. Thus, we reasoned that it may act as a dietary restriction mimetic and reduce mTORC1 activity. We tested whether acarbose inhibited mTORC1 signaling in the brains of *Ndufs4^-/-^* mice by looking at the phosphorylation of ribosomal protein S6 (S6RP) via Western blot. As expected, rapamycin ablated phosphorylation of S6RP in both wild type and *Ndufs4^-/-^* animals. Conversely, mice treated with acarbose showed no appreciable reduction in S6RP phosphorylation (Figure 3A, B).

**Figure 3:**
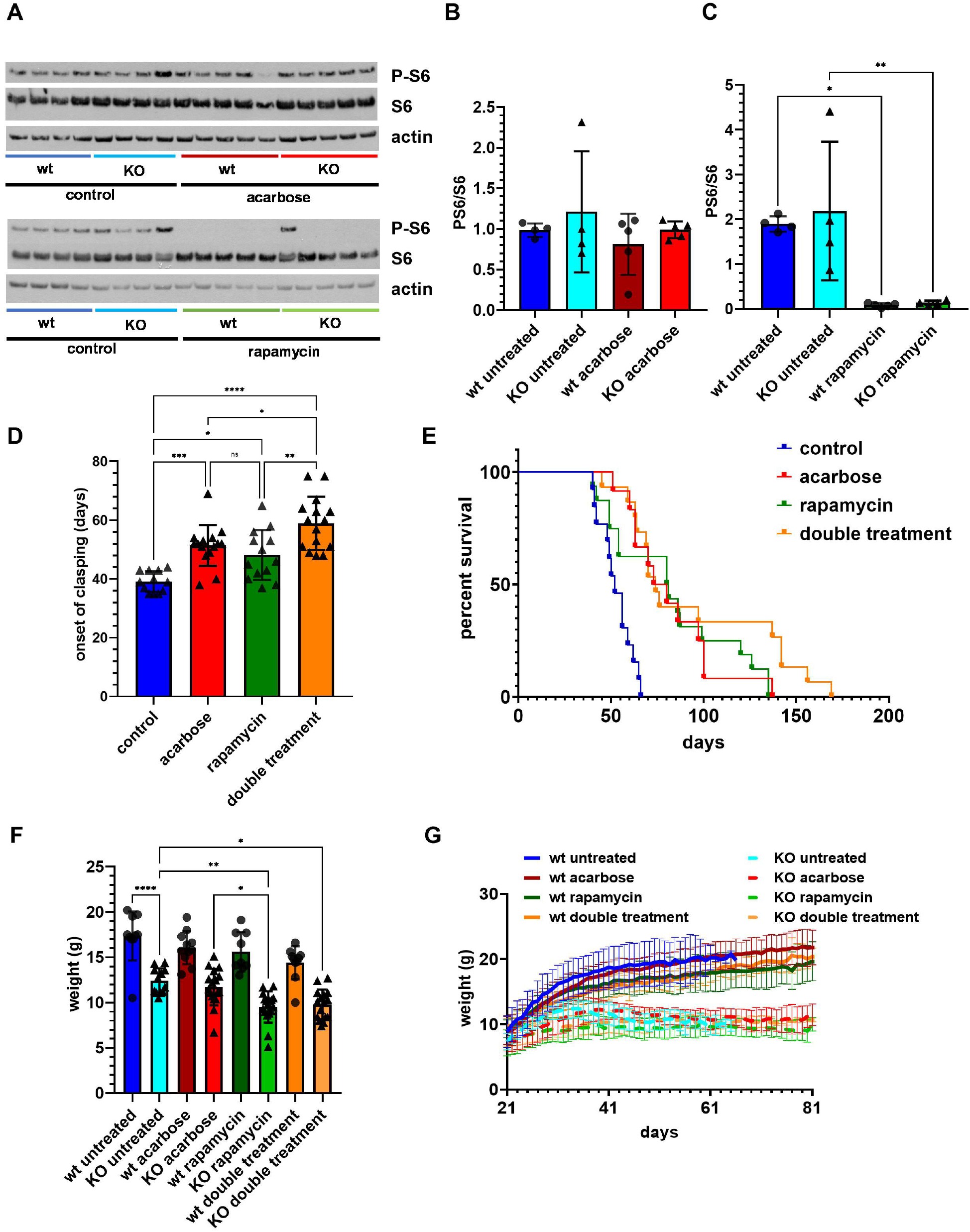
Acarbose does not inhibit mTORc1 signaling but does not further extend lifespan in *Ndufs4^-/-^* mice treated with rapamycin. **A.** Representative western blots of ribosomal protein S6 phosphorylation at serine 235/236. **B. C.** Densitometric analysis of the western blots in panel A. *<0.05, **: p<0.01 one way ANOVA. N=4 wild type and *Ndufs4^-/-^* untreated, N=5 wild type and *Ndufs4^-/-^* acarbose or rapamycin treated **D.** Onset of neurological symptoms (clasping) measured in days after birth in control-chow (blue,) 0.1% acarbose-chow (red) fed, 8mg/kg/day I.P. rapamycin-treated (green,) acarbose-chow fed and every other day rapamycin treated (orange) *Ndufs4^-/-^* mice. *: t-test p <0.05, **: p < 0.01 one-way ANOVA. N=12 *Ndufs4^-/-^* untreated, N=15 *Ndufs4^-/-^* acarbose, N=13 *Ndufs4^-/-^* rapamycin, N=15 *Ndufs4^-/-^* double treated **E.** Survival curves of *Ndufs4^-/-^* mice fed either control (blue,) 0.1% acarbose diet (red,) treated with 8 mg/kg daily intraperitoneal rapamycin (orange,) acarbose diet + every other day rapamycin (orange). Median lifespan was 52 days for control-chow fed, 76.5 days for acarbose-chow fed mice (log-rank p<0.05), 80 for rapamycin treated mice (log-rank p<0.05), and 74 for double treatment regimens (log-rank p<0.01). Double treatment with acarbose and every other day rapamycin also increased maximum lifespan, Mann-Whitney U, p<0.05. N=13 *Ndufs4^-/-^* untreated, N=12 *Ndufs4^-/-^*acarbose, N=16 *Ndufs4^-/-^* rapamycin, N=15 *Ndufs4^-/-^* double treated **F.** Comparison of body weight progression at 35 days post-natal for untreated wt (dark blue), untreated *Ndufs4^-/-^* (light blue), acarbose treated wt (dark red), acarbose treated *Ndufs4^-/-^* (light red), rapamycin treated wt (dark green), rapamycin treated *Ndufs4^-/-^* (light green), double treated wt (dark orange), and double treated *Ndufs4^-/-^* mice (light orange). *: p<0.05. **: p<0.01, ***: p<0.001, ****: p<0.0001, n.s.: not significant, one-way ANOVA. N=10 wild type untreated, N=13 *Ndufs4^-/-^* untreated, N=11 wild type acarbose-treated, N=17 *Ndufs4^-/-^*acarbose-treated, N=9 wild type rapamycin-treated, N=16 *Ndufs4^-/-^* rapamycin-treated, N=10 wild type double treated, N=15 *Ndufs4^-/-^* double treated. **G.** Weight progression from weaning until post-natal day 81. Solid dark blue: wild type mice fed control chow. Solid dark red: wild type mice fed acarbose chow. Dotted light blue: *Ndufs4^-/-^* mice fed control chow. Dotted light red: *Ndufs4^-/-^* mice fed acarbose chow. Solid dark green: wt rapamycin. Dotted light green: *Ndufs4^-/-^* rapamycin. Solid dark orange: wt double treated. Dotted light orange: *Ndufs4^-/-^* double treated. N=10 wild type untreated, N=13 *Ndufs4^-/-^* untreated, N=11 wild type acarbose-treated, N=17 *Ndufs4^-/-^*acarbose-treated, N=9 wild type rapamycin-treated, N=16 *Ndufs4^-/-^* rapamycin-treated, N=10 wild type double treated, N=15 *Ndufs4^-/-^* double treated.

To further assess the interaction between rapamycin and acarbose, we tested whether administering both rapamycin and acarbose would further delay the onset of neurological phenotypes and improve survival in *Ndufs4^-/-^* mice. We administered 1000 ppm acarbose in the chow and daily intraperitoneal injections of 8 mg/kg rapamycin to *Ndufs4^-/-^* mice and control littermates beginning at weaning (post-natal day 21). To our surprise, neither control nor KO mice tolerated the double treatment regimen. The mice were unable to gain weight (Figure S3) and died within a week. To avoid this lethal effect, we reduced the frequency of rapamycin injection to every other day. As expected, acarbose delayed the onset of clasping by 10 days. Interestingly, the effects of daily rapamycin when treatment is initiated at weaning were comparable to those of acarbose, and less than when treatment is initiated at post-natal day 10 ^7^. Clasping was further delayed by another 10 days in mice receiving both acarbose and every other day rapamycin, indicating that the two drugs have additive effects on the suppression of neurological symptoms (Figure 3C). Next, we determined whether double treatment with both acarbose and rapamycin could further increase survival in *Ndufs4^-/-^* mice compared to either treatment alone. Surprisingly, all three drug regimens (acarbose, daily rapamycin, acarbose and every other day rapamycin) had similar effects on median lifespan extension (Figure 3D). However, mice receiving both acarbose and every other day rapamycin showed a significant increase in maximum survival compared to either untreated or acarbose-treated animals. Importantly though, the double treatment did not further reduce weight and growth progression in *Ndufs4^-/-^* mice compared to rapamycin alone (Figure 3F and G).

### Acarbose reverses metabolic dysfunction and prevents the accumulation of glycolytic intermediates in the olfactory bulb of *Ndufs4^-/-^* mice

*Ndufs4^-/-^* mice display prominent neurometabolic defects once neurological symptoms emerge, including accumulation of glycolytic intermediates and reduction of both fatty and amino acids in brain and liver ^7^. Since acarbose significantly delays the appearance of inflammatory markers in the regions most affected by the lack of *Ndufs4*, we performed region-specific targeted metabolic profiling of pre-symptomatic (post-natal day 30) *Ndufs4^-/-^* mice fed either control or acarbose chow in order to determine if 1) metabolic alterations precede, and may thus be causal to, onset of disease and brain pathology, and 2) whether acarbose can prevent these alterations and generally rescue metabolic defects in these animals. We detected 138 metabolites across 4 different regions of the mouse brain via LC-MS (Figure 4A). Principal Component Analysis (PCA) showed a clear separation by genotype in the olfactory bulb when comparing the metabolic profile of untreated wild type and *Ndufs4^-/-^* animals, and separation by treatment when comparing untreated *Ndufs4^-/-^* animals with *Ndufs4^-/-^* animals treated with acarbose (Figure S4A). We employed a linear model fitting to test the group differences within each brain region. As expected from the PCA analysis, the olfactory bulb showed the greatest number of significantly altered metabolites (47) between wild type and *Ndufs4^-/-^* animals. Consistent with previous reports ^7^, *Ndufs4^-/-^* mice showed an accumulation of early glycolytic intermediates (Figure 4B). Treatment with acarbose significantly restored wild type levels of most of these metabolites in the olfactory bulb (Figure 4C). Next, we performed metabolite set enrichment (MSEA) and pathway analysis (MPA) to determine which metabolic pathways were significantly altered in the olfactory bulb of *Ndufs4^-/-^* mice prior to onset of disease. MSEA revealed that the first 9 pathways differentially regulated between control and acarbose-treated *Ndufs4^-/-^* mice are all involved in carbohydrate metabolism, with glycolysis and gluconeogenesis as the most enriched (Figure S4B). Similarly, carbohydrate metabolic pathways were prominently featured in MPA, together with glutathione metabolism and branched-chain amino acids biosynthesis (Figure S4C). We mapped glycolytic intermediates onto glycolysis via specific pathway analysis to better understand where the changes in glycolysis occur between untreated and untreated mice (Figure 4D, E.) Specific pathway analysis showed that early glycolytic intermediates were significantly higher in the olfactory bulb of *Ndufs4^-/-^* mice, and acarbose specifically lowered their levels to those of wild type brains and significantly reduced pyruvate levels, further confirming that impairment in glycolysis is an early feature of disease in *Ndufs4^-/-^* mice, and suggesting that preventing this impairment may recapitulate the effects of acarbose.

**Figure 4:**
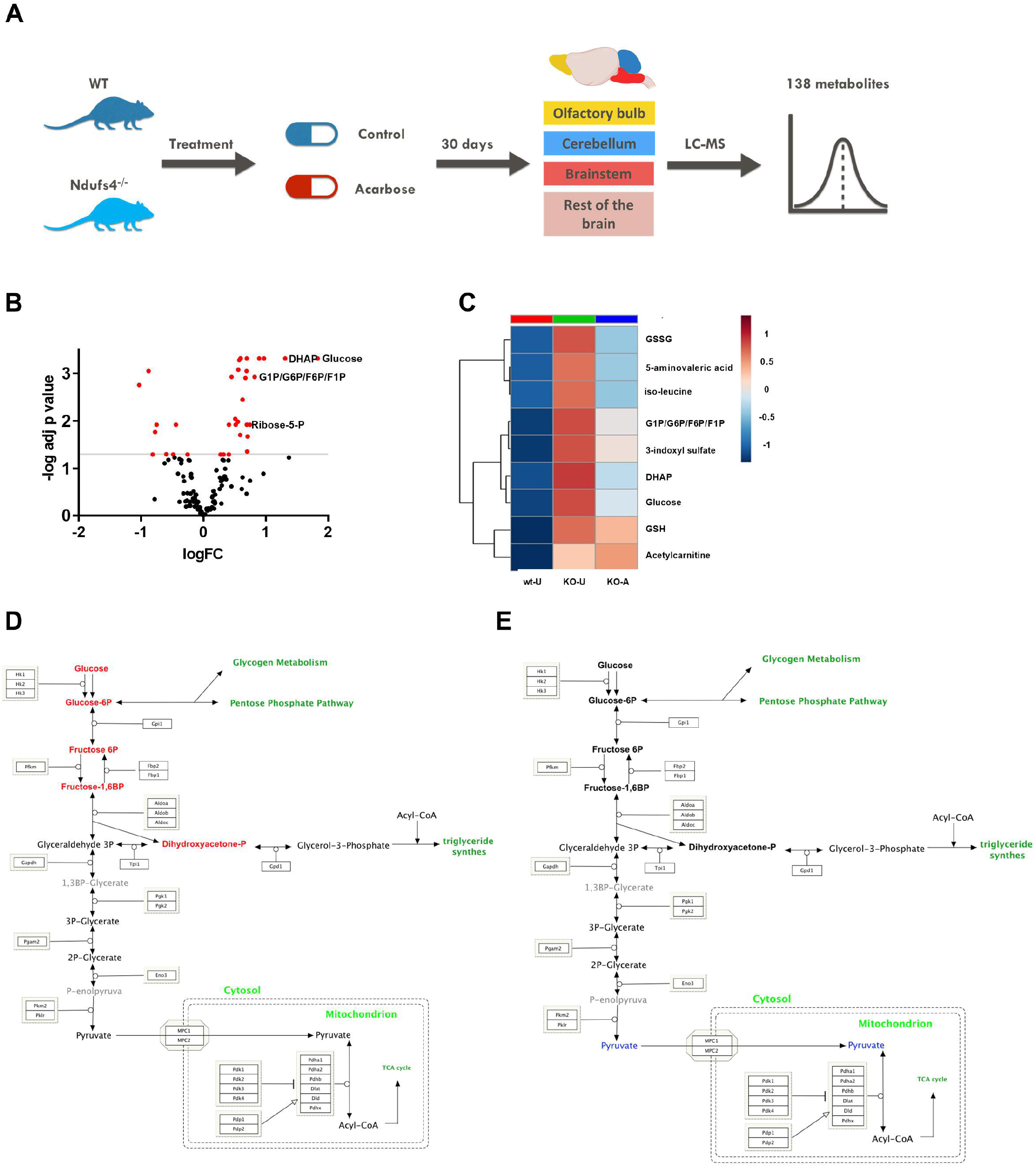
Metabolic profiling of brain regions in *Ndufs4^-/-^* mice treated with acarbose. **A.** Schematic diagram of the metabolomics assay. **B.** Volcano plot of metabolic features in the olfactory bulb of untreated *Ndufs4^-/-^* mice compared to wild type littermates. Red dots are significantly altered metabolites. **C.** Heat map of the relative abundance of selected significantly altered metabolites in the olfactory bulb of wild type untreated (WT-U), *Ndufs4^-/-^* untreated (KO-U), and acarbose-treated *Ndufs4^-/-^* mice (KO-A). **D.** and **E.** Specific Pathway Analysis of glycolysis intermediates in the olfactory bulb of untreated (D.) and acarbose-treated (E.) *Ndufs4^-/-^* mice. Red metabolites are significantly more abundant compared to untreated wild type; blue metabolites are significantly less abundant. N=4-5 per group.

We hypothesized that acarbose may relieve the overload of glycolytic intermediates by reducing glucose uptake and thus the rate of glycolysis. We compared 1-hour post-prandial levels of circulating glucose in wild type and *Ndufs4^-/-^* mice treated with acarbose with those of untreated animals by measuring blood glucose concentration one hour after the beginning of the dark cycle, when mice tend to consume their biggest meal of the day. As expected, acarbose reduced post-prandial glucose in wild type animals. However, acarbose did not further reduce blood glucose concentration in *Ndufs4^-/-^* mice, which already had lower levels than wild type mice (Figure S4D). Next, we sought to determine whether direct inhibition of glycolysis could relieve symptoms of disease, similar to acarbose treatment.

To test this hypothesis, we treated *Ndufs4^-/-^* mice with two hexokinase inhibitors, 2-deoxyglucose (2-DG) and glucosamine (GlcN). These drugs are reported to reduce glycolytic flux accumulation of intermediates. We monitored onset of clasping and survival. Surprisingly, neither drug increased survival nor delayed symptoms of disease in *Ndufs4^-/-^* mice (Figure S4F, E).

### Acarbose remodels the cecal microbiome in *Ndufs4^-/-^* mice and alters the concentration of short chain fatty acids

While conducting necropsies on *Ndufs4^-/-^* mice treated with acarbose, we noticed that their ceca and large intestines were significantly larger than those of untreated mice (Figure 5A.) Cecum hypertrophy is a common feature of germ-free, gnotobiotic, and acarbose-fed rodents that is associated with remodeling of the intestinal flora ^57–60^. In order to determine whether acarbose alters the intestinal flora of *Ndufs4^-/-^* mice, we performed 16S rDNA sequencing of the cecal content of *Ndufs4^-/-^* and wt littermates fed either a control diet or a diet supplemented with acarbose. Linear regression analysis of operational taxonomic units at the genus level identified a significant reduction in the levels of Bacteroides (p=7.3×10^-5^, FDR=0.001). Analysis also showed nominal significance (p<0.05, FDR=0.14) for reduction in Prevotella and increased abundance of Clostridium, Rikenella, and Alistipes in the colons of acarbose-treated animals (Figure 1B, 1C, Supplementary Figure 5 A-D). The effects of acarbose on murine lifespan have been linked to an increased abundance of anaerobic enteric bacteria and production of short-chain fatty acids ^61^. Bacterially derived short chain fatty acids improve nutrient metabolism and mitochondrial function in multiple organs, including the brain, by increasing mitochondrial thermogenesis and energy expenditure, providing substrates for beta-oxidation and gluconeogenesis, and modulating the activity of enzymes involved in glycolysis, TCA cycle, and fatty acid catabolism in general ^62,63^. We measured the concentration of 12 SCFAs in the cecum of untreated and acarbose-treated mice via LC-MS ^64^ and obtained absolute concentrations for acetic, propionic, iso-butyric, butyric, 2-methyl-butyric, iso-valeric, valeric, and caproic acids (table 1). Pairwise comparison of SCFAs concentrations revealed that acarbose significantly decreased the concentration of acetic, iso-butyric, 2-methyl-butyric, iso-valeric, and valeric acid (Figure 5 D, F, H, I, J), while the levels of butyric acid were significantly increased (Figure 5 G). Propionic and caproic acid levels were unaffected by acarbose (Figure 5 E, K).

**Figure 5:**
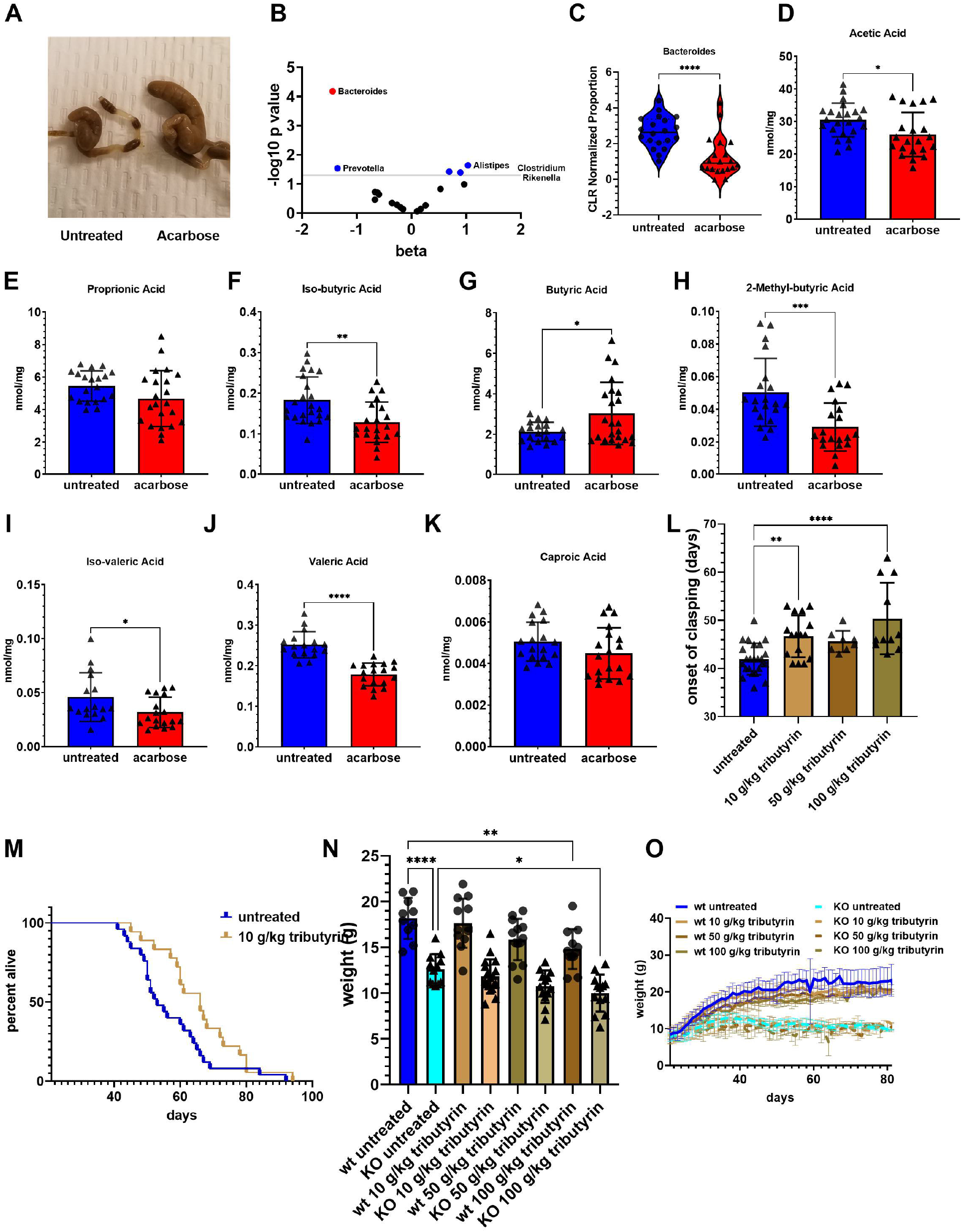
Acarbose remodels the microbiome and its metabolism in *Ndufs4^-/-^* mice. **A.** Representative ceca of untreated (left) and acarbose-treated (right) *Ndufs4^-/-^* mice. **B.** Volcano plot of bacterial genera in the ceca of acarbose-treated mice compared to untreated animals. Blue dots are nominally significant (p<0.05 linear regression analysis) for altered abundance by acarbose treatment. Red dots are significantly altered by acarbose treatment with false discovery rate, FDR<0.05. **C.** Centered log-ratio normalized proportion of Bacteroides in the ceca of untreated and acarbose-treated animals. ****: p<0.0001 Student’s t test. **D-K.** Concentration of acetic (D.) propionic (E.) iso-butyric (F.) butyric (G.) 2-methyl-butyric (H.) iso-valeric (I.) valeric (J.) and caproic (K.) acid in the ceca of untreated and acarbose-treated mice. *: p<0.05. **: p<0.01, ***: p<0.001, ****: p<0.0001 Student’s t test. N= 20/group panels A to K **L.** Onset of clasping in *Ndufs4^-/-^* treated with increasing doses of tributyrin. N=24 untreated, N= 15 10 g/kg tributyrin, N=8 50 g/kg tributyrin, N=11 100 g/kg tributyrin. **M.** Survival plot of *Ndufs4^-/-^* treated with 10 g/kg tributyrin. Log-rank p<0.05. N=25 untreated, N=18 10 g/kg tributyrin **N.** Comparison of weights at 35 days post-natal for control-chow fed wild type, *Ndufs4^-/-^* mice, and tributyrin fed mice. *: p<0.05. **: p<0.01, ***: p<0.001, ****: p<0.0001 one way ANOVA. N=10 wild type untreated, N=12 *Ndufs4^-/-^* untreated, N=12 wild type 10 g/kg tributyrin, N=18 *Ndufs4^-/-^* 10 g/kg tributyrin, N=13 wild type and *Ndufs4^-/-^* 50 g/kg tributyrin, N=12 wild type 100 g/kg tributyrin, N=15 100 g/kg tributyrin. **O.** Weight progression from weaning until post-natal day 81. Solid dark blue: wild type mice fed control chow (dark blue), *Ndufs4^-/-^* untreated (light blue), and tributyrin (shades of brown). N=13 wild type and *Ndufs4^-/-^* untreated, N=12 wild type 10 g/kg tributyrin, N=18 *Ndufs4^-/-^* 10 g/kg tributyrin, N=13 wild type and *Ndufs4^-/-^* 50 g/kg tributyrin, N=12 wild type 100 g/kg tributyrin, N=15 100 g/kg tributyrin.

**Table 1:** Short Chain Fatty Acids monitored via LC-MS and their average detection level in mouse ceca.

| SPQC SCFA | Average (nmol/mg) | CV % |
| --- | --- | --- |
| Acetic acid | 2.73E+01 | 0.44% |
| Propionic acid | 4.93E+00 | 1.70% |
| iso-butyric acid | 1.63E-01 | 1.89% |
| butyric acid | 2.58E+00 | 1.75% |
| 2-methyl butyric acid | 5.03E-02 | 0.54% |
| iso-valeric acid | 5.07E-02 | 2.00% |
| valeric acid | 2.15E-01 | 0.92% |
| 3-methyl valeric acid | BLOQ | BLOQ |
| iso-caproic acid | BLOQ | BLOQ |
| caproic acid | 4.96E-03 | 9.56% |
| Heptanoic acid | BLOQ | BLOQ |
| Octanoic acid | BLOQ | BLOQ |

### Butyric acid supplementation recapitulates some of the effects of acarbose

Since acarbose significantly alters the concentration of several SCFAs in the cecum of *Ndufs4^-/-^* mice, we hypothesized that manipulating dietary SCFAs might recapitulate some of its effect on disease progression and survival. Butyric acid was more represented in the ceca of both our animals and HET3 mice treated with acarbose ^61^. Thus, we asked whether butyric acid supplementation could alleviate disease progression. To test this hypothesis, we treated *Ndufs4^-/-^* mice with increasing doses of tributyrin, a triacyl-glycerol ester of butyric acid, which was more represented in the ceca of our animals as well as HET3 mice treated with acarbose ^61^. Increasing doses of tributyrin delayed onset of clasping, with 100 g/kg having effects comparable to those of acarbose (Figure 5 L.) The lowest dose of tributyrin tested (10 g/kg) significantly extended median lifespan in *Ndufs4^-/-^* mice by 25%, while 50 g/kg and 100 g/kg extended survival by 13% (Figure 5M, Figure S5E). Increasing doses of tributyrin also negatively affected weight gain in *Ndufs4^-/-^* mice as evinced by a decreasing linear trend in both wt and *Ndufs4^-/-^* animals and significantly lower weights in animals treated with 100 g/kg (Figure 5 N,O, Figure S5F, G).

## Discussion

The data presented here demonstrate that acarbose is an effective treatment for mitochondrial disease in a mouse model of Leigh Syndrome (Figures 1 and 2). We tested acarbose in this context based solely on its previously documented ability to slow aging in wild type mice ^51,52^, as a test of the hypothesis that aging and mitochondrial disease share common underlying mechanisms. To the best of our knowledge, acarbose or other alpha-glucosidase inhibitors have never been tested or proposed as a treatment for mitochondrial disease. These data are consistent with the geroscience approach to health, which posits that many, if not all, chronic age-related diseases share biological aging as a common underlying mechanism ^65,66^. Our results suggest that the geroscience approach can be successfully applied to diseases that are not obviously age-related, like childhood mitochondrial disorders. Acarbose and rapamycin now represent two distinct examples supporting this idea, with further support provided by independent work showing that other geroprotective interventions including hypoxia ^67,68^, α-ketoglutarate ^69^, and nicotinamide mononucleotide ^70^ can enhance survival in the *Ndufs4^-/-^* model ^71–73^.

The mechanism of action for acarbose in the *Ndufs4^-/-^* mice is distinct from that of rapamycin, at least at the level of mTOR inhibition. Rapamycin is a specific and potent inhibitor of mTORC1 whereas acarbose has no detectable effect on mTORC1 activity in tissues, and combination treatment with acarbose and rapamycin has partial additive effects for both neurological symptoms and survival in the *Ndufs4^-/-^* mice. Interestingly, rapamycin and acarbose are each able to suppress the aberrant accumulation of glycolytic intermediates in *Ndufs4^-/-^* brain, suggesting that they may impinge on overlapping downstream mechanisms of disease rescue. It will be interesting to see whether co-administration of acarbose and rapamycin will similarly improve age-related outcomes in wild type mice, relative to acarbose or rapamycin alone. We unexpectedly observed synthetic lethality in very young mice treated with daily rapamycin at 8 mg/kg combined with 1000 ppm acarbose in the chow. This lethality was not associated with loss of *Ndufs4*, as both knockout and wild type mice were equally affected. Reducing the rapamycin treatment to every-other-day suppressed the effect as did beginning double treatment in older mice (not shown,) suggesting that the synthetic lethality results from strong mTOR inhibition combined with acarbose treatment during early development.

Metabolic imbalances arise early in some brain regions of *Ndufs4^-/-^* mice and could be causal to the disease rather than a consequence of pathology, since they are reversed by acarbose treatment before the appearance of symptoms (Figure 4.) However, treatment with glycolytic inhibitors failed to suppress symptoms of disease in this model. Although we haven’t directly tested the efficacy of 2-deoxyglucose and glucosamine to inhibit glycolysis in *Ndufs4^-/-^* mice, the doses we tested have been well characterized and shown to reduce circulating glucose, improve insulin sensitivity, and inhibit glycolysis in mouse ^74,75^. It is unclear whether a similar metabolic profile would arise later in life in the olfactory bulb of acarbose treated mice, in association with the onset of neurological phenotypes, as we have not performed region-specific metabolomics on older animals. However, acarbose still reduces glial activation in the brain at post-natal day 50 (Figure 1,) which would suggest that protection from metabolic alterations may be maintained in these animals. The tissues analyzed for this study were also used to analyze the effects of rapamycin on metabolite changes in specific brain regions ^76^. Notably, two different methods of analysis yielded similar metabolic profiles in the olfactory bulb of untreated *Ndufs4^-/-^* mice, further strengthening the validity of our results.

Rather than directly modulating systemic metabolism, we propose that acarbose is acting, at least in part, by altering the composition of the intestinal microbiome and therefore the production of secondary metabolites in the large intestine. Indeed, acarbose treated mice show a marked reduction in the abundance of Bacteroides and Prevotella, while the abundance of Alistipes, Clostridia, and Rikenella was increased (Figure 5). These results are consistent with observations in patients with type 2 diabetes treated with acarbose, where drug treatment consistently reduced the abundance of Bacteroides and efficacy of treatment correlated positively with baseline Bacteroides counts ^77^. In aging wild type mice, acarbose has been shown to increase the production of most short-chain fatty acids through intestinal anaerobic bacteria ^61^. However, in our study we find that the production of several SCFAs is reduced by acarbose treatment, while production of butyric acid is increased. These differences may be ascribed to the different nature of the samples analyzed (feces vs cecal content), the strain (HET3 vs C57BL6), the vivarium location, or a combination of these factors. Indeed, a more detailed and carefully controlled analysis of the intestinal microbiome both in the context of aging and mitochondrial disease is necessary to understand the beneficial effects of acarbose in both conditions. Intriguingly, the relative abundance and metabolic activity of Bacteroides seems to be influenced by the luminal concentration of butyric acid, which can lower the pH of proximal areas of the large intestine, including the cecum ^78^. We found that supplementation with a butyric acid source, tributyrin, was sufficient to partially phenocopy disease rescue by acarbose. While we were unable to obtain the same magnitude of effect from tributyrin as from acarbose, we note that butyrate is only one of several bioactive fermentation products altered by acarbose administration (Figure 5). Additional strategies to modulate the levels of several SCFAs may prove more effective than perturbing the levels of only one molecule.

In conclusion, we have identified a potential new avenue of treatment for Leigh Syndrome and other mitochondrial diseases with acarbose, an already FDA-approved anti-diabetic drug that is generally considered to be safe ^56^ or through dietary supplementation with SCFAs. The proposed mechanism of action holds the potential to advance treatment of these disorders, as short-chain fatty acids and probiotic supplements are easily obtained and administered even to pediatric patients. While additional studies are needed to validate this hypothesis and the effectiveness of alpha-glucosidase inhibitors in other mitochondrial disorders, the current study establishes a foundation on which new approaches can be built.

## Acknowledgements

We thank Carly Horne, Ayush Sharma, Adrian Markewych, Thomas Krivak, Alvine Nguouonga, Kelly Wang, Alex Naini, and Noah Biru for additional help with animal husbandry and treatment. We thank the husbandry and veterinary staff of the Foege/ARCF facilities at the University of Washington for their assistance in caring for the animals.

## Author Contributions

A.B. and M.K. devised the study. A.B., A.S.G., B.M.G.N, H.T., K.S., N.T., A.M., administered treatments, collected weight data, and scored neurological phenotypes and survival. A.B., H.T., K.S., and N.T., collected tissues and ran western blot analyses. A.B., A.S.G., B.M.G.N., H.T., K.S., N.T., A.M., S.R.U., helped maintain the mouse colony and genotyped animals, I.B.S. and R.W.D. helped with the design, execution, and analysis of microbiome sequencing data, K.Y. and L.W. helped with the analysis of metabolomics data, E.B.K. helped with the design and collection of samples for metabolomics analysis, J.M.S. performed all histopathology scoring, D.L.S Jr. contributed acarbose and scientific rationale for the study, J.W.T. and L.D. measured the abundance of SCFAs, A.B. and M.K. wrote the manuscript.

---

**Figure S1:**
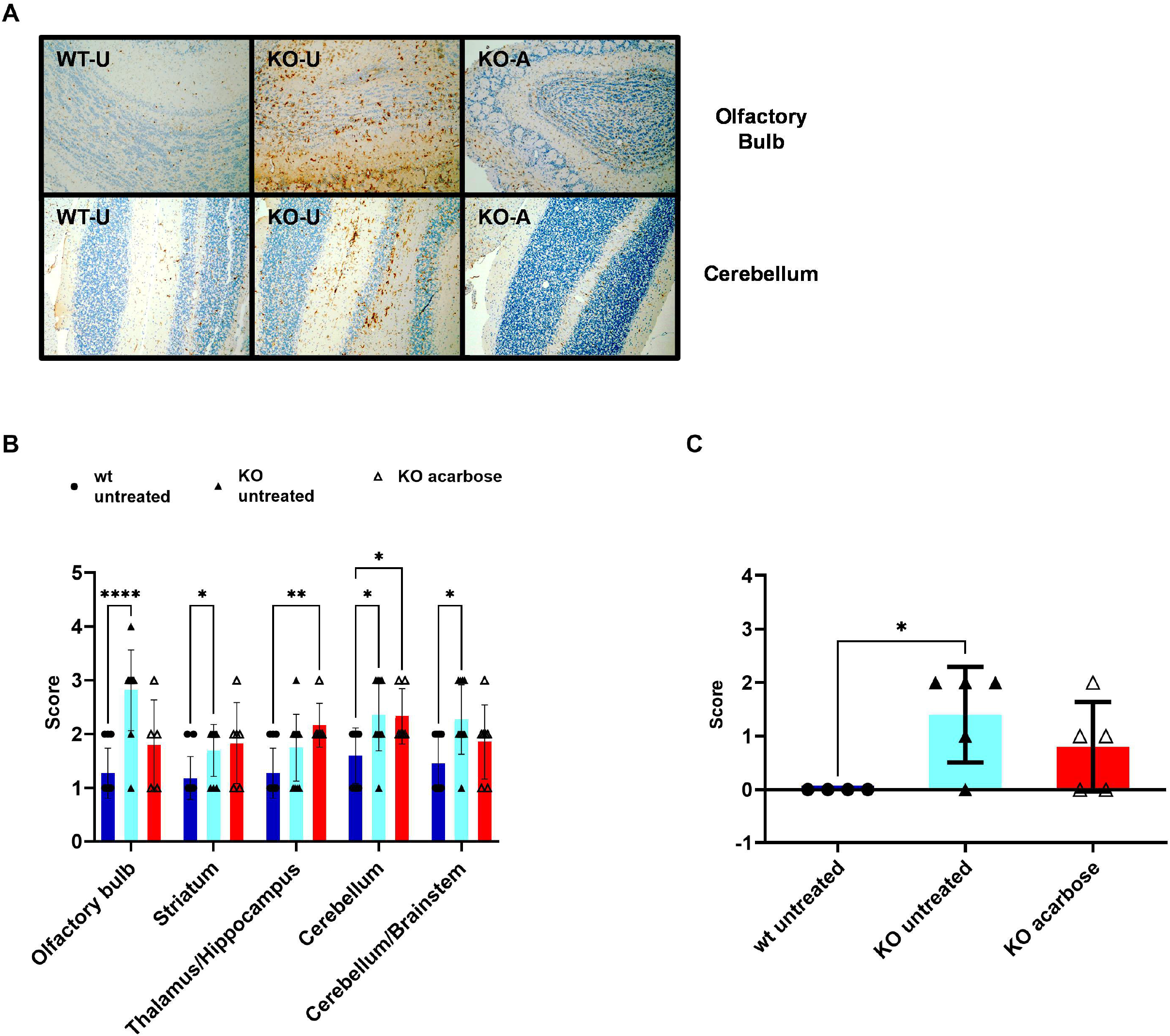
Acarbose reduces brain lesions in *Ndufs4^-/-^* mice. **A.** Representative images of Iba1 immunostaining in the olfactory bulb and cerebellum. **B.** Intensity of Iba1 immunostaining in the sections described in panel A **: p<0.01 *: p<0.05, mixed effects analysis. **C.** Vacuolation score in the vestibular nuclei of wild type, *Ndufs4^-/-^*, and acarbose-chow fed *Ndufs4^-/-^* mice. Slides were scored on a 0-4 scale with 0=absent and 4=severe vacuolation. *: p<0.05 one-way ANOVA. N=4-5/group

**Figure S2.**
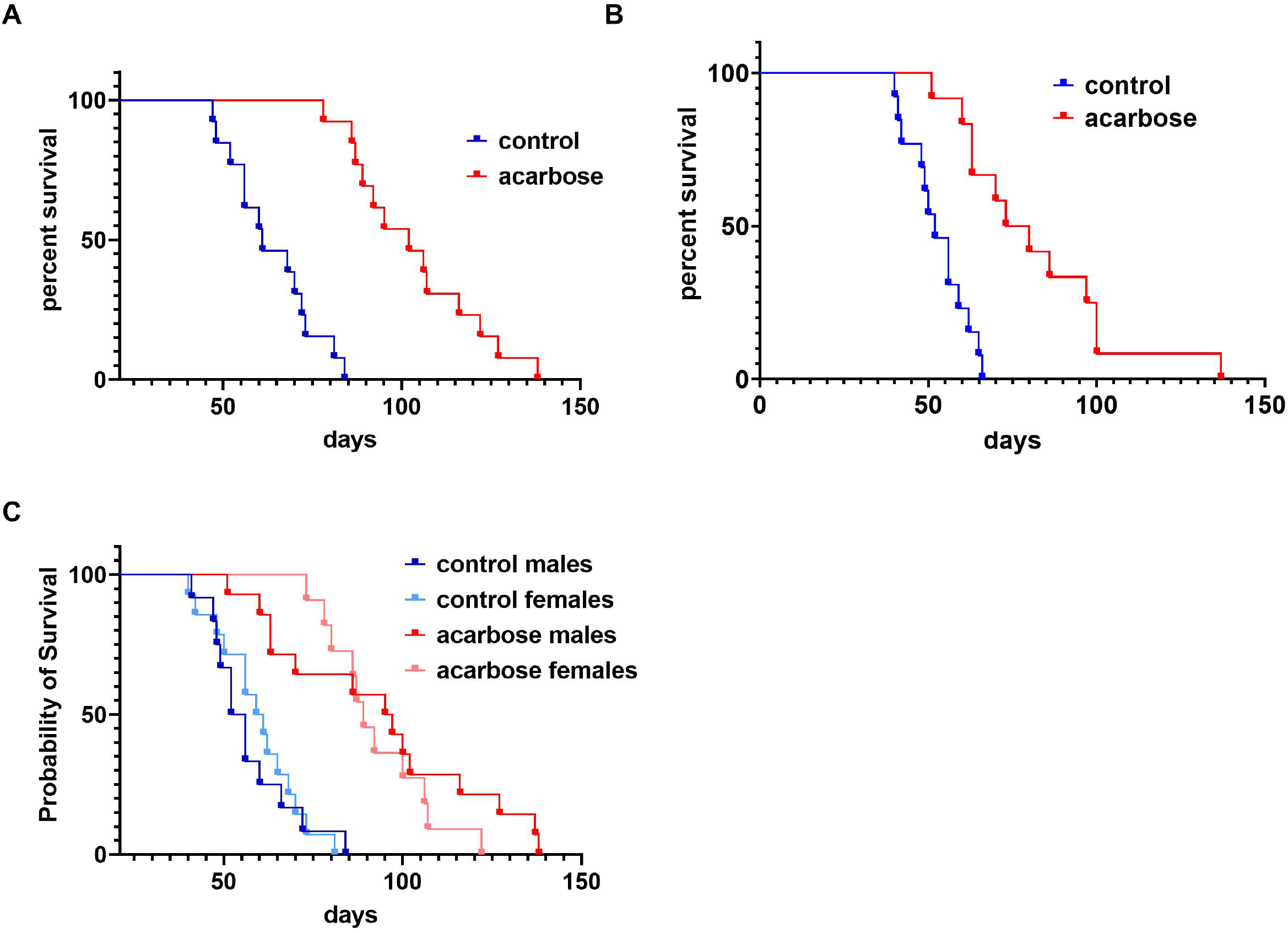
A. and B. Individual survival curves of control-chow and acarbose-chow fed *Ndufs4^-/-^* mice. A. Log-rank p<0.0001 N=13/group B. Log-rank p<0.001 N=13 untreated, N=12 acarbose-treated C. Pooled lifespan curves divided by sex. Log-rank p<0.5944, N=14 acarbose-treated males, N=11 acarbose-treated females.

**Figure S3:**
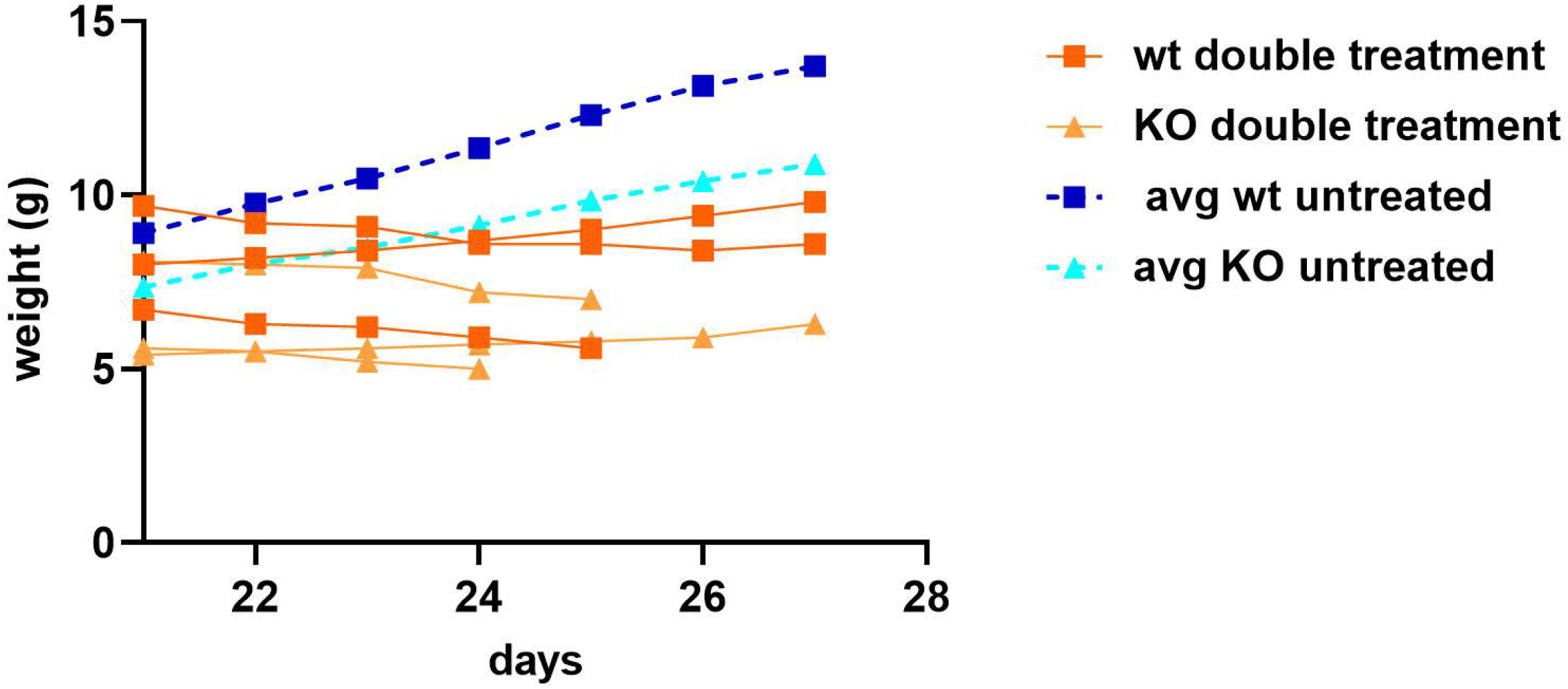
Daily double treatment with acarbose and rapamycin prevents growth and ultimately causes death in newly weaned mice. Solid lines indicate weight progression individual animals on daily double treatment with rapamycin and acarbose from weaning. Dark orange squares: wild type animals, light orange triangles: *Ndufs4^-/-^* animals Dotted lines are average weights of untreated animals, dark blue squares: wild type untreated, light blue triangles, *Ndufs4^-/-^* untreated.

**Figure S4:**
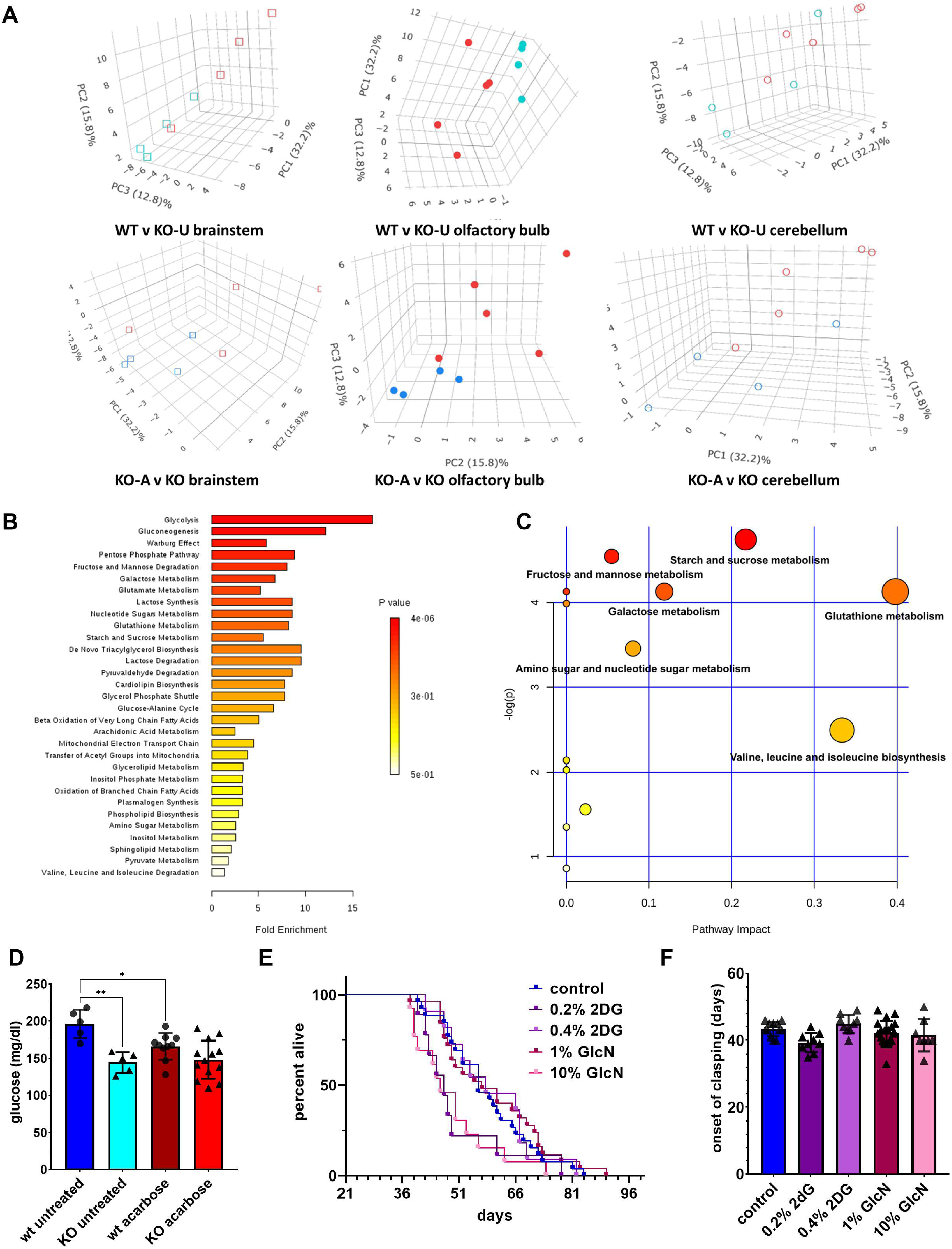
**A.** Principal component analysis of the metabolomics dataset described in figure 4A. Light blue squares and circles, wild type untreated; red squares and circles, *Ndufs4^-/-^* untreated; dark blue squares and circles, acarbose-treated *Ndufs4^-/-^*. **B.** Metabolites Set Enrichment Analysis (MSEA) of metabolic pathways enriched in the olfactory bulb of untreated *Ndufs4^-/-^* compared to wild type animals. **C.** Metabolites Pathway Analysis (MPA) of metabolic pathways enriched in the olfactory bulb of untreated *Ndufs4^-/-^* compared to wild type animals. Color denotes p-value, same scale as in panel B., size of circles denotes impact. N=4-5 per group for panels A to C. **D.** 1h post-prandial blood glucose levels in wild type and *Ndufs4^-/-^* mice untreated or treated with acarbose. N=5 wild type and *Ndufs4^-/-^* untreated, N=10 wild type N=13 *Ndufs4^-/-^* acarbose-treated. **E.** Survival plot of *Ndufs4^-/-^* mice treated with 0.2%, 0.4% 2-deoxyglucose (2DG) or 1%, 10% glucosamine (GlcN) mixed in the food. N=26 untreated, N=9 0.2% 2-deoxyglucose, N=11 0.4% 2-deoxyglucose, N=25 1% glucosamine, N=13 10% glucosamine. **F.** Onset of clasping in *Ndufs4^-/-^* mice treated with 2-deoxyglucose (2DG) or glucosamine (GlcN). N=12 untreated, N=9 0.2% and 0.4% deoxyglucose, N=21 1% glucosamine, N=8 10% glucosamine.

**Figure S5:**
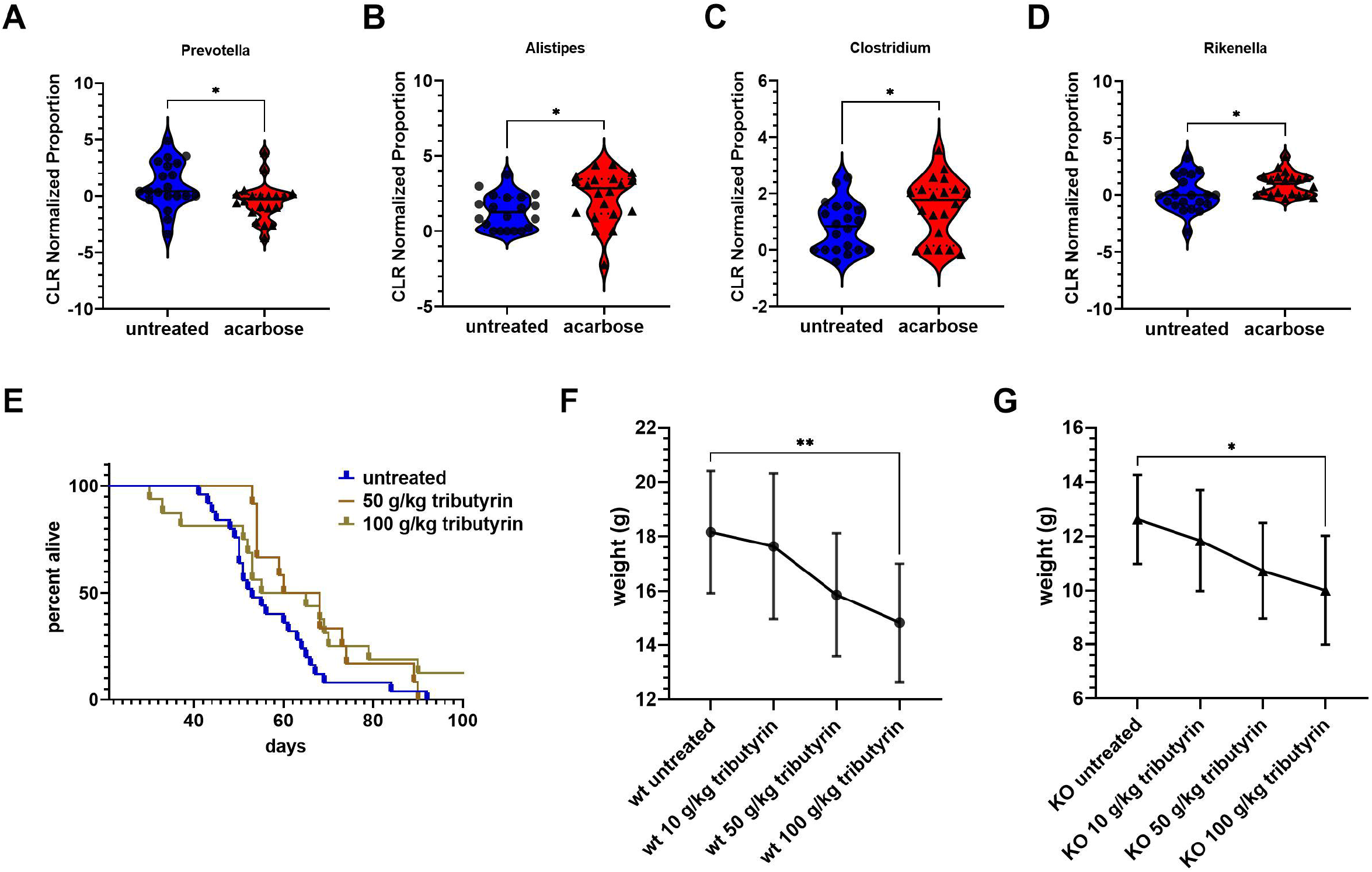
**A-D.** Centered log-ratio normalized proportion of Prevotella (A.) Alistipes (B.) Clostridium (C.) and Rikenella (D.) *: p<0.05 Student’s t test. N= 20/group **E.** Survival plot of *Ndufs4^-/-^* treated with 50mg/kg or 100 g/kg tributyrin. N=25 untreated, N=12 50 g/kg tributyrin, N=16 100 g/kg tributyrin **F-G.** Linear trend of wt (F.) and *Ndufs4^-/-^* mice (G.) weight after treatment with increasing doses of tributyrin at post-natal day 35. P<0.001 one-way ANOVA test for linear trend. N=13 wild type and *Ndufs4^-/-^* untreated, N=12 wild type 10 g/kg tributyrin, N=18 *Ndufs4^-/-^* 10 g/kg tributyrin, N=13 wild type and *Ndufs4^-/-^* 50 g/kg tributyrin, N=12 wild type 100 g/kg tributyrin, N=15 100 g/kg tributyrin.

## RESOURCE AVAILABILITY

### Lead Contact

Further information and requests for resources and reagents should be directed to and will be fulfilled by the lead contact, Matt Kaeberlein

### Materials Availability

This study did not generate new unique reagents

### Data and Code Availability

All data reported in this paper will be shared by the lead contact upon request.

This paper does not report original code.

Any additional information required to reanalyze the data reported in this paper is available from the lead contact upon request.

## EXPERIMENTAL MODEL AND SUBJECT DETAILS

### Mice

C57BL6/N *Ndufs4^-/-^* mice and *Ndufs4^+/+^* littermates were generated by breeding heterozygous pairs in the University of Washington Foege/ARCF animal vivarium. All animals were genotyped prior to post-natal day 21 and littermate pairs of different genotypes were randomly assigned to treatments beginning around post-natal day 21. Mice that weighed less than 6.5 grams at postnatal day 21 were kept in the breeder cage until they reached that weight or post-natal day 28, whichever occurred first. For 16S sequencing, wild type and *Ndufs4^-/-^* mice from different breeder pairs were housed together to control for parental effects on microbiome composition. All animals were maintained in the Foege/ARCF animal vivarium at the University of Washington and housed in groups of 2-5 in either Allentown JAG 75 or Allentown NexGen Mouse 500 cages, in temperature-controlled rooms (25° C) on racks providing filtered air and filtered, acidified water. All animals were housed on a 14h light, 10h dark cycle. Animals were fed Pico Lab Diet 20 5053 (St. Louis MO) and treated with 1000 ppm acarbose (Tecoland, Irvine CA) mixed in the chow, 10 mg/kg, 50 mg/kg, or 100 mg/kg tributyrin (Fisher Scientific, Waltham, MA) mixed in the chow, 1% or 10% glucosamine (Fisher Scientific), 0.2% or 0.4% 2-deoxyglucose (Millipore-Sigma, Billerica CA), or daily intraperitoneal injections of 8 mg/kg rapamycin (LC laboratories, Woburn MA). For survival curves and monitoring of neurological symptoms, animals were monitored 2-3 times a week until onset of symptoms and then monitored daily until endpoint. Animals were euthanized when they showed any of the following signs: 1) loss of over 30% of the maximum weight recorded, 2) inability to eat or drink, 3) severe lethargy, as indicated by a lack of response such as a reluctance to move when gently prodded, 4) severe respiratory difficulty while at rest, indicated by a regular pattern of deep abdominal excursions or gasping. Animals utilized for western blotting, metabolomics, 16S sequencing, and SCFAs measurements were fasted overnight and refed for 3h at the beginning of the following day’s light cycle before cervical dislocation. Tissues were snap frozen in liquid nitrogen and stored under liquid nitrogen for further analysis. Animals utilized for histology were anesthetized with a ketamine/xylazine mix and perfused with ice cold PBS followed by 10% neutral buffered formalin prior to dissection.

### Histology

Following perfusion, brains were removed from the skull and immersion fixed in 10% neutral buffered formalin. Coronal sections of brain at the level of the olfactory lobe, striatum, thalamus with hippocampus, and two sections of cerebellum/medulla were obtained and routinely paraffin embedded, sectioned at 4-5 microns, and stained with hematoxylin and eosin (HE) by the Histology and Imaging Core are the University of Washington. Immunohistochemistry for Iba1 and GFAP were performed by Harborview Medical Center Pathology Lab. Anti-GFAP (Agilent, Dako Omnis, M0761) was used at a 1:300 dilution. Anti Iba1 (Wako Chemicals, 019-19741) was used at a 1:500 dilution. Slides were processed with Leica Bond III IHC stainer, using Heat Induced Epitope retrieval with Bond Epitope Retrieval Solution 1 for 20 minutes, and Bond Polymer Refine detection kit (Leica Biosystems, Buffalo Grove IL). H&E and immunohistochemistry slides were scored blindly by a board-certified veterinary pathologist (J.M.S.). Vacuolation was scored on a 0-4 scale with 0=normal; 1=minimal; 2=mild; 3=moderate and 4=severe lesions. Intensity of immunohistochemical staining was scored semi-quantitatively on a 1-4 scale with 1 indicating expected baseline staining; 2 indicating a focal cluster of more strongly immunopositive cells or multifocal or diffuse mildly increased staining; 3 indicating moderately increased staining (multifocal or multifocal to coalescing) and 4 indicating markedly increased coalescing to diffuse staining).

### Western Blotting

Frozen tissues were ground in liquid nitrogen and protein were extracted in ice-cold RIPA buffer supplemented with protease and phosphatase inhibitors. 30 μg of protein extract were loaded onto NuPage Bis-Tris acrylamide gels (Thermofisher, Waltham MA), and transferred onto Immobilon P membranes (Millipore-Sigma, Burlington MA) with a BioRad Trans-Blot Turbo Transfer System (BioRad, Hercules CA). Membranes were blotted with antibodies against phospho-S6 (Ser 235–236, cat 2211), total S6 (clone 5G10) (Cell Signaling Technologies, Danvers MA).

### Metabolomics

The olfactory bulb, brainstem, and cerebellum were dissected from freshly isolated brains and snap frozen for further analysis. All samples were submitted to the Northwest Metabolomics Research Center at the University of Washington.

Brain samples were thawed at 4° C and homogenized in 200 uL PBS. 800 μL of methanol containing 53.1 μM ^13^C-arginine and 50.1 μM ^13^C-glucose (Sigma-Aldrich, Saint Louis, MO) was added, and the samples were then incubated for 30 min at −70 °C. The suspension was sonicated in an ice bath for 10 min and then centrifuged at 20,800 *g* for 10 min at 0–4 °C. Supernatants (600 μL) were collected and dried for 60 min using an Eppendorf Vacufuge Drier (Happauge, NY), then each reconstituted in 600 μL 40% Solution A and 60% Solution B, also containing 5.13 μM ^13^C-tyrosine and 22. 6 μM ^13^C-lactate. Solution A was 30 mM ammonium acetate in 85% H_2_O/ 15% acetonitrile and 0.2% acetic acid. Solution B was 15% H_2_O/ 85% acetonitrile and 0.2% acetic acid. Reagents for these solutions were purchased from Fisher Scientific (Pittsburgh, PA). All samples were then filtered through Millipore PVDF filters (Pittsburgh, PA) immediately prior to chromatography.

The LC system was composed of two Agilent 1260 binary pumps, an Agilent 1260 auto-sampler and Agilent 1290 column compartment containing a column-switching valve (Agilent Technologies, Santa Clara, CA). Each sample was injected twice, 10 μL for analysis using negative ionization mode and 2 μL for analysis using positive ionization mode. Both chromatographic separations were performed in HILIC mode on two parallel Waters XBridge BEH Amide columns (150 × 2.1 mm, 2.5 μm particle size, Waters Corporation, Milford, MA). While one column was performing the separation, the other column was reconditioned and ready for the next injection. The flow rate was 0.300 mL/min, the auto-sampler temperature was kept at 4 °C, and the column compartment was set at 40 °C, and total separation time for both ionization modes was 20 min. The gradient conditions for both separations were identical and consisted of 10% Solvent A from 0 to 2 min, a ramp to 50% during min 2–5, continued 50% Solvent A from 5 to 9 min, a ramp back to 10% from 9 to 11 min, and then 10% Solvent A from 11 to 20 min.

After the chromatographic separation, MS ionization and data acquisition was performed using an AB Sciex QTrap 5500 mass spectrometer (AB Sciex, Toronto, ON, Canada) equipped with an electrospray ionization (ESI) source. The instrument was controlled by Analyst 1.5 software (AB Sciex, Toronto, ON, Canada). Targeted data acquisition was performed in multiple-reaction-monitoring (MRM) mode. The source and collision gas was N2 (99.999% purity). The ion source conditions in negative mode were: Curtain Gas (CUR) = 25 psi, Collision Gas (CAD) = high, Ion Spray Voltage (IS) = - 3.8KV, Temperature (TEM) = 500 °C, Ion Source Gas 1 (GS1) = 50 psi and Ion Source Gas 2 (GS2) = 40 psi. The ion source conditions in positive mode were: CUR = 25 psi, CAD = high, IS = 3.8KV, TEM = 500 °C, GS1 = 50 psi and GS2 = 40 psi.

### Blood Glucose Measurements

Blood glucose was measured one hour after the beginning of the dark cycle with a Alpha Trak 2 glucometer at day 35 post-natal (Zoetis, Parsipanny NJ).

### 16S rDNA sequencing

Cecum content was collected at the time of necropsy and snap frozen in liquid nitrogen. Samples were submitted to RTL Genomics (Lubbock, TX) for sequencing and identification. rDNA extracts were sequenced on Illumina MySeq sequencers. Raw sequences were denoised using the following steps: 1) forward and reverse reads in FASTQ format were merged together using the PEAR Illumina paired-end read merger, 2) Paired FASTQ files were then converted into FASTA formatted sequences and trimmed back at the last base with a total average read greater than 25. 3) Sequences were sorted by length and run through the USEARCH algorithm to perform prefix dereplication and clustering at 4% divergence. 4) Operational Taxonomic Units were selected with the UPARSE OTU algorithm and chimera checking was performed on the selected OTUs with the UCHIME detection software in *de novo* mode. 5) Clusters were then mapped to their corresponding OTUs, chimeric sequences removed, and individual reads mapped to the clusters. OTUs taxonomic assignment were determined by USEARCH alignments against a database of high-quality sequences derived from the NCBI database.

### Short Chain Fatty Acid Analysis

50-100 mg of cryoground cecum content was weighed into Precellys Soft Tissue Homogenizing CK14 tubes (Bertin, Rockville MD) and shipped to the Duke Proteomics and Metabolomics Core at Duke University for analysis. Samples were extracted and derivatized 3-nitrophenylhydrazine as previously described ^64^. Briefly, 50/50 ethanol water was added at a ratio of 10 uL per 1 mg. All tubes were homogenized on the Precellys 24 bead blaster (Bertin Instruments, Rockville MD) at 10° C for 3 cycles at 10,000 rpm. 20 uL from each sample was derivatized by adding 20 uL of 200 mM 3-nitrophenyl hydrazine (3-NPH) in 50% ethanol with 6% pyridine and 120 mM N-(3-dimethylaminopropyl)-N’-ethylcarbodiimide hydrochloride (EDC·HCl) in 50% ethanol, incubating at 35oC for 30 minutes while shaking. The reaction was quenched with 760 uL of cold 10% ethanol in water with 1% formic acid and held at 4° C. Data collection was performed using LC-MS/MS on a Sciex 6500+ QTRAP mass spectrometer (Sciex, Framingham MA), including calibration curves for each analyte and a ^13^C6 internal standard for each compound (via ^13^C6 NPH derivatization reagent).

UPLC separation of the SCFAs was performed using a Exion AD liquid chromatograph (Sciex, Framingham, MA) with a Waters Acquity 2.1 mm x 50 mm 1.7 um BEH C18 column fitted with a Waters Acquity C18 1.7 um Vanguard guard column. Analytes were separated using a gradient from 15% solvent B to 100% solvent B in 9.5 minutes. Solvents A & B were 0.1% formic acid in water and acetonitrile, respectively. The total UPLC analysis time was approximately 13.5 minutes. The method uses electrospray ionization in negative mode introduced into a 6500+ QTrap mass spectrometer (Sciex) operating in the Multiple Reaction Monitoring (MRM) mode. MRM transitions (compound-specific precursor to product ion transitions) for each analyte and internal standard were collected over the appropriate retention time window. The standard curve was run once at the beginning of the run queue and once at the end of the sample run queue. The Golden West QCs, study pool QC (SPQC) and calibration QCs were run once at the beginning, once in the middle and once at the end of the run queue. Injection volume for each injection was 5 uL.

All data was analyzed in Skyline v21.1.1 (www.skyline.ms) ^79^ which includes raw data import, peak integration, and a linear regression fit with 1/x2 weighting for the calibration curves.

The calibration curves for all the analytes contain 12 points, from 0.98 uM to 10 mM for AA, and from 0.098 to 1 mM for the other analytes. Each calibrator’s residual bias was calculated after regression fit, and any point other than the lower and upper limits of quantitation where the residual fell outside of 15% were removed from the calibration curve equation. The lower limit of quantitation (LLOQ) was defined to be the lowest point that meets the criteria of having a bias < 20% and the upper limit of quantitation (ULOQ) is defined in the same way for the highest point. The calibration range that was used for concentration calculations in the samples is listed in Table 1 for each analyte.

### Statistical Analysis

Unless otherwise stated, all data were analyzed using the GraphPad Prism 9 software. P values of 0.05 or lower was considered statistically significant.

Statistical analysis for metabolomics data was carried out using R (version 4.0.2). The targeted metabolomics assay was designed to detect 210 metabolites and 138 of these metabolites were detected in at least one samples. We performed a median normalization where we adjusted the data, so all samples have the same median value of the metabolite abundance post log_2_ transformation. 135 metabolites with < 20% missingness and a coefficient of variation (CV) < 20% in the pooled sample QC data were included in further analysis. After filtering, no missing values remained, therefore, no imputation was performed. Principal Component Analysis was performed on the transformed and normalized dataset using R prcomp() function. We fit linear models to the normalized and filtered data using the Bioconductor limma package ^80^ to assess the difference in abundance between the groups within each brain region while adjusting for the sample collection month in the model. The limma package uses empirical Bayes moderated statistics, which improves power by ‘borrowing strength’ between metabolites in order to moderate the residual variance ^81^. We selected metabolites with a false discovery rate (FDR) of 10%Pathway analysis were conducted with MetaboAnalyst 5.0 ^82^.

### 16S Sequence Taxonomic Regression Analysis

The proportional 16S read abundances of each sample’s genus level taxa were centered log ratio transformed using the clr() function of the ‘compositions” R package (https://cran.r-project.org/web/packages/compositions/index.html). Linear regression models using the base R lm() function were used to determine the acarbose exposure treatment association with the clr transformed taxa outcome while adjusting for sex and cage segregation. Common taxa with a mean proportional abundance >0.01 were tested. False discovery rate (FDR) adjustment to p-values were made using the R p.adjust() function with the BH (Benjamini and Hochberg) method ^83^. Nominal significance was set at p<0.05 and statistically significant taxa are reported with an FDR<0.05.

## Funding

3 P30 AG 013280 Nathan Shock Center of Excellence for the Biology of Aging

1 R01 NS98329 Mechanisms of Mitochondrial Disease Suppression in Ndufs4 Knockout Mice

